# Engineering microalgal cell wall-anchored proteins using GP1 PPSPX motifs and releasing with intein-mediated fusion

**DOI:** 10.1101/2025.01.23.634604

**Authors:** Kalisa Kang, Évellin do Espirito Santo, Crisandra Jade Diaz, Stephen Mayfield, João Vitor Dutra Molino

**Affiliations:** Department of Molecular Biology, School of Biological Sciences, University of California San Diego, La Jolla, CA, United States of America; Department of Biochemical and Pharmaceutical Technology, Faculty of Pharmaceutical Sciences, University of São Paulo, São Paulo, Brazil; Algenesis Inc., 1238 Sea Village Dr., Cardiff, CA, United States of America

**Keywords:** microalgae, cell wall anchoring, glycomodules, pH-sensitive intein, intein-mediated cleavage

## Abstract

Harnessing and controlling the localization of recombinant proteins is essential for advancing synthetic biology, industrial biotechnology, and drug delivery. This study presents a dual system for protein anchoring and controlled release in *Chlamydomonas reinhardtii*. Using truncated variants of the GP1 glycoprotein fused to the plastic-degrading enzyme PHL7, we identified the PPSPX motif as critical for anchoring proteins to the cell wall. Constructs with increased PPSPX content exhibited reduced secretion but enhanced anchoring, revealing key anchor-signal sites within GP1. To enable controlled release, we incorporated a pH-sensitive intein from derived from *Mycobacterium tuberculosis* RecA. Under acidic conditions (pH 5.5), this intein efficiently cleaved and released mCherry and PHL7 from the GP1 anchor. Fluorescence kinetics demonstrated significant mCherry release from the mCherry-intein-GP1 construct within 4 hours at pH 5.5, while release was minimal at pH 8.0 and negligible for the mCherry-GP1 control. Western blot analysis confirmed efficient cleavage, showing a lower band corresponding to free mCherry at pH 5.5 and no release at pH 8.0. This anchor-release strategy integrates glycomodules with pH-sensitive inteins to enable precise spatial and temporal protein control. The system offers broad utility for targeted drug delivery, environmental biosensing, and biocatalyst deployment. Overall, we establish a versatile framework for optimizing protein localization and environmentally triggered release in *C. reinhardtii*, with broad implications for proteomics, biofilm engineering, and scalable therapeutic delivery systems.

**Graphical Abstract:** 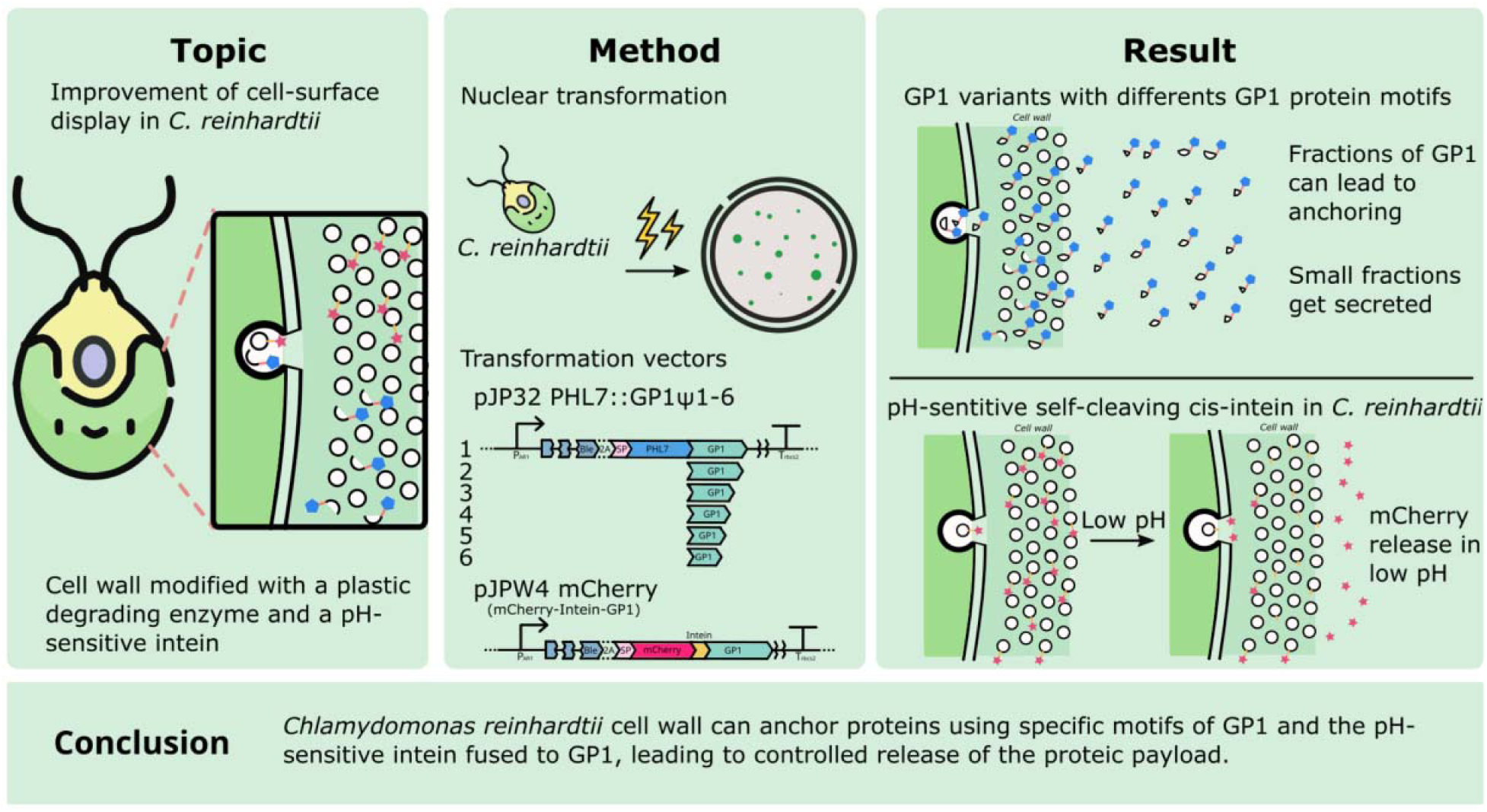

## 1. Introduction

The ability to precisely control protein localization, anchoring, and release is crucial for a wide range of biotechnological applications, including industrial bioprocessing, drug delivery, and gene therapy [1–4]. As the field advances, there is an increased demand for systems that can both stably anchor proteins and conditionally release them under specific stimuli or conditions. Microalgae have emerged as a promising platform for engineering these complex biological systems. Their robust cell walls, ease of genetic manipulation, and potential for large-scale cultivation make them attractive candidates for applications ranging from biofuel production to the expression of recombinant proteins [5]. However, challenges remain in achieving precise control over protein localization and release in these organisms, which is essential for optimizing their use in various industrial and therapeutic applications [6,7].

One promising approach to address these challenges involves the use of glycomodules: short peptide sequences rich in serine-proline (SP) repeats and proline-proline-serine-proline-X (PPSPX) repeats derived from native microalgal cell wall proteins, such as the hydroxyproline-rich GP1 protein attached to the cell wall via non-covalent interactions [8–10]. The existing literature conveys that when synthetic (SP)_n_ glycomodules were attached to the C-terminus of proteins of interest expressed in *Chlamydomonas reinhardtii* and *Nicotiana tabacum*, protein secretion was enhanced [11–13]. Increased secretion is a feature that is highly desirable for downstream purification and applications requiring high protein yields. The capacity to enhance protein secretion has significant implications for drug delivery systems. Proteins designed for therapeutic use often require high yield and purity to be cost-effective, particularly when produced at scale. By integrating glycomodules into the design of therapeutic proteins, their secretion-enhancing properties can be exploited to produce biologics, such as monoclonal antibodies, growth factors, or enzyme therapies, in microalgal platforms [14]. However, contrary to these results, this *C. reinhardtii* study demonstrated that when a plastic-degrading enzyme, PHL7, [15] was fused to GP1-derived glycomodules containing PPSPX motifs, PHL7 was anchored to the cell wall rather than secreted.

Increased secretion or stable anchoring are advantageous, but there are also many biotechnological applications that require a mechanism for the controlled release of anchored proteins [7,16]. To address the challenge of controlled release of proteins, the use of inteins was explored. Inteins, or intervening proteins, are autocatalytic protein elements that posttranslationally splice or cleave themselves from a host protein to form the mature protein [17–19]. Self-splicing or -cleavage of these protein elements can be engineered to be induced by changes in light, pH, temperature, or redox state, and to be responsive to the addition of small molecules [17–19]. Splicing or cleavage by inteins does not require energy supply, exogenous host-specific proteases, or cofactors [18,20]. Genetically engineering microalgae to anchor therapeutics or enzymes to the cell wall that could be released upon specific stimuli offers a scalable and cost-effective alternative to other drug delivery systems, such as micro or nanorobots [21,22]. Additionally, combining the stable anchoring provided by microalgae with the precision of microrobot technology could enhance drug delivery by merging cost-effectiveness with precise targeting. Similarly, in industrial bioprocessing, the controlled release of enzymes or functional proteins at specific stages of the process can significantly enhance both efficiency and yield. For instance, the use of thermostable inteins to control the activity of cell wall-degrading enzymes, such as xylanases in genetically engineered maize, has demonstrated the potential to avoid detrimental effects on plant growth while still enabling high enzyme activity during processing [23]. Applying a similar strategy in microalgae could allow for the controlled release of enzymes at optimal stages in bioprocessing, improving the overall efficiency and scalability of biofuel and biochemical production. This study uses a cis-cleaving, pH-sensitive intein that self-cleaves at its C-terminal at pH 5.5[24]. By integrating this pH intein into the vector constructs, a system was developed to not only anchor proteins to the cell wall, but also allow for their controlled release in response to a change in pH.

This study combines glycomodules and pH-sensitive inteins to create a versatile platform for targeted protein localization and release in microalgae. The anchoring capacity of glycomodules was validated using PHL7, demonstrating its effective localization to the cell wall. Subsequently, vectors that fused GP1 and PHL7 or mCherry via the pH-sensitive intein (PHL7-intein-GP1 or mCherry-intein-GP1, respectively) were engineered, transformed into *C. reinhardtii*, and assessed for their ability to mediate controlled protein release under acidic conditions. The results showed that, upon acid treatment, the inteins cleaved as designed, releasing the downstream proteins into the supernatant. By providing a system that can anchor proteins with high specificity and release them under controlled conditions, this study opens up new possibilities for the development of advanced therapeutic delivery systems, more efficient industrial bioprocesses, and other applications in biotechnology.

## 2. Results

### 2.1 Glycomodule-mediated anchoring to *C. reinhardtii*’s cell wall

To investigate whether the truncated portions of GP1 can serve as anchoring domains for proteins on the *C. reinhardtii* cell wall, we constructed and analyzed six recombinant vectors expressing PHL7 fused to truncated variants (glycomodules) of GP1. The glycomodules were generated by amplifying the GP1 coding region from the mCherry-GP1 vector [10] via PCR. Due to the high GC content and repetitive structure of the GP1 sequence, truncated variants were produced. This process yielded six sequences, each containing a distinct number of motif repeats (**Figure 1**). These truncated GP1 variants were fused to the C-terminus of PHL7, resulting in the recombinant constructs designated as pJP32PHL7_X, where X corresponds to 0p1s, 2p16s, 3p20s, 13p20s, 15p20s, or 18p20s. The naming scheme denotes the number of specific motifs: “p” represents the number of PPSPX motifs, and “s” indicates the number of SP motifs. For example, the 0p1s variant contains one SP motif but no PPSPX motifs, whereas the 18p20s variant contains 18 PPSPX motifs and 20 SP motifs. The pJP32PHL7 construct serves as the baseline strain expressing secreted PHL7 without GP1 fusion.

**Figure 1:**
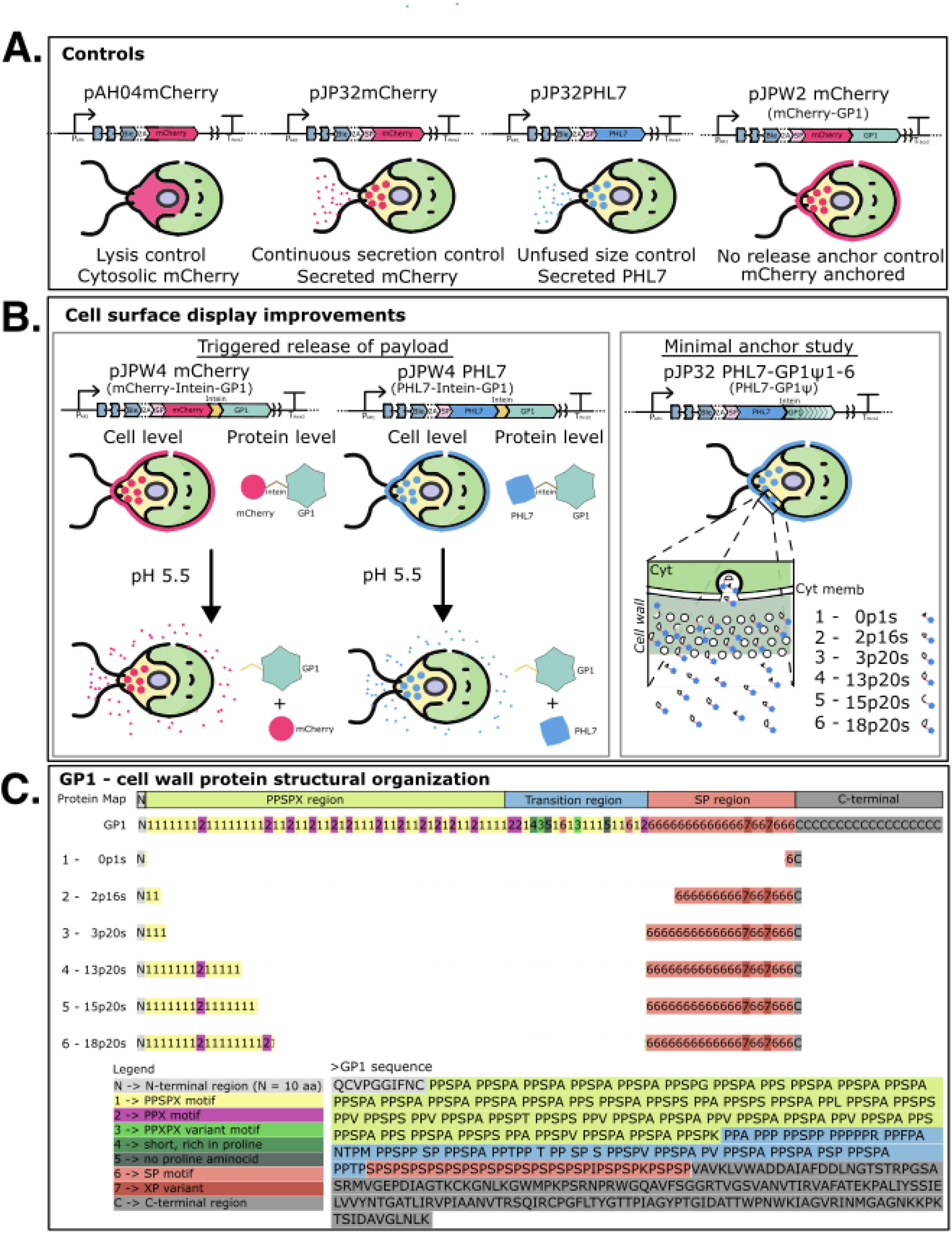
Schematic of cell surface display improvements. **(A)** The control strains used in this study are pAH04mCherry (cytosolic mCherry), pJP32mCherry (secreted mCherry), pJP32PHL7 (secreted PHL7), pJPW2mCherry (mCherry-GP1; cell wall-anchored mCherry). **(B)** We explored 2 strategies to improve cell surface display. The first involves the induced release of a payload via a cis-cleaving pH intein at pH 5.5. The pJPW4mCherry construct contains an mCherry-intein-GP1 fused protein that releases mCherry from the cell wall into the extracellular environment upon pH 5.5 buffer treatment. Similarly, the pJPW4PHL7 construct contains an PHL7-intein-GP1 fused protein that releases PHL7 upon pH 5.5 buffer treatment. The second strategy involves anchoring PHL7 onto the cell wall via glycomodules of varying lengths that contain different motifs. **(C)** The structural organization of GP1 from *C. reinhardtii* and the glycomodule constructs used in the second strategy to improve cell surface display are described here. The N-terminal region (labeled ‘N’ in gray) spans the first 10 amino acids of the mature form. The PPSPX region (shaded green) is enriched in repetitive PPSPX motifs (labeled “1”) and PPX motifs (“2”). Following this, the transition region (shaded blue) is characterized by increased sequence variability, including with some PPSPX variant motifs (“3”) exhibiting slight sequence divergence, short proline-rich segments (“4”), PPSPX motifs, and SP motifs (“6”). The SP region (red) contains repetitive SP motifs and terminates in the C-terminal region (gray), a more typical non-repetitive sequence that includes three predicted N-glycosylation sites, marked as “C”. Recombinant GP1 variants (0p1s, 2p16s, 3p20s, 13p20s, 15p20s, and 18p20s) were designed by truncating the C-terminal region and redistributing motifs from the PPSPX and SP regions. Each variant retains portions of the N-terminal and SP regions, with differing quantities of PPSPX and SP motifs. The naming convention for the variants reflects the number of motifs they contain, with “p” indicating the number of PPSPX motifs and “s” denoting the number of SP motifs. The primary amino acid sequence of GP1 is shown below the map, with each functional region highlighted for clarity.

The pJP32PHL7_X vectors were transformed into *C. reinhardtii* via electroporation, and the cells were plated onto TAP agar plates containing 15 μg/mL zeocin and 0.75% (v/v) Impranil® DLN. Every transformation generated colonies with clearing zones, or halos, in the opaque polyester polyurethane polymer suspension that becomes transparent upon degradation by PHL7 (**Table 1**) [25]. The vectors containing more PPSPX motifs (13p20s, 15p20s, and 18p20s) had a lower proportion of colonies with halos compared to that of vectors containing less PPSPX motifs (0p1s, 2p16s, and 3p20s). This suggests that the PPSPX motif may be involved with anchoring rather than secretion [11–13].

**Table 1:** Quantification of colony formation and halo production by *C. reinhardtii* strains expressing various vector constructs. The table summarizes the total number of colonies*, the number of colonies displaying halos, and the calculated proportion of colonies with halos for each vector construct. *The total colonies refers to the total number of colonies for each transformation experiment.

| Vector construct | Total colonies | Colonies with halos | Proportion of colonies with halos |
| --- | --- | --- | --- |
| pJP32PHL7_0p1s | 1551<br>1545 | 442<br>262 | 0.285<br>0.170 |
| pJP32PHL7_2p16s | 1393<br>1403 | 252<br>161 | 0.181<br>0.115 |
| pJP32PHL7_3p20s | 989<br>1689 | 81<br>499 | 0.082<br>0.295 |
| pJP32PHL7_13p20s | 1488<br>1296 | 2<br>2 | 0.0013<br>0.0015 |
| pJP32PHL7_15p20s | 1290<br>1198 | 6<br>4 | 0.0047<br>0.0033 |
| pJP32PHL7_18p20s | 1124<br>1130 | 10<br>7 | 0.0089<br>0.0062 |
| pJP32PHL7_control | 1502<br>1387 | 343<br>363 | 0.228<br>0.262 |

The colonies with halos were picked into 96-well plates, along with the pJP32PHL7 control and wild-type, and grown for 7 days. Constructs pJP32PHL7_13p20s, 15p20s, and 18p20s generated significantly less halos than the control (ANOVA followed by Dunnett’s multiple comparison test; adjusted p-values = 0.0354, 0.0372, and 0.0397, respectively). An Impranil® DLN-degrading activity assay was employed to further screen for PHL7 activity. Impranil® DLN, an opaque polyester polyurethane polymer suspension, becomes transparent upon degradation. Since PHL7 cleaves ester bonds in polymers [15,25], transformants secreting PHL7 formed transparent halos around the colonies on the opaque plates, indicating both secretion and activity of PHL7. This halo phenotype was leveraged to measure the decrease in absorbance as the enzyme degraded the polymer. The activity assay revealed a negative trend: as the number of motifs repeats increased, there was a decrease in activity (i.e., change OD at 350 nm over 2 days [day 5 to day 7]) (**Figure 2A**). We observed significantly lower activity in each glycomodule vector compared to the pJP32PHL7 control (ANOVA followed by Tukey post-hoc analysis; adjusted p-values < 0.0001; n = 84 for every vector). The pJP32PHL7 baseline strain is used as a benchmark for soluble, unconstrained enzyme activity in this assay. Additionally, there was a statistically significant difference between the 2p16s and 3p20s vectors (adjusted p-value 0.0122). On the other hand, the 13p20s, 15p20s, and 18p20s vectors displayed no activity (with the exception of an outlier for 15p20s), with levels comparable to the wild type for the supernatant samples (**Figure 2A**). To further confirm the presence of each truncated glycomodule in the vectors, the transformants exhibiting the highest activity levels were selected and their supernatants were used to run a zymogram containing 1% (v/v) Impranil® DLN. As the number of motifs repeats increased, the band size of clearing zones also increased (**Figure 2C**). For pJP32PHL7_control and 0p1s, two bands were observed, conveying that two active isoforms of the enzyme were present due to glycosylations sites present in PHL7 [25].

**Figure 2:**
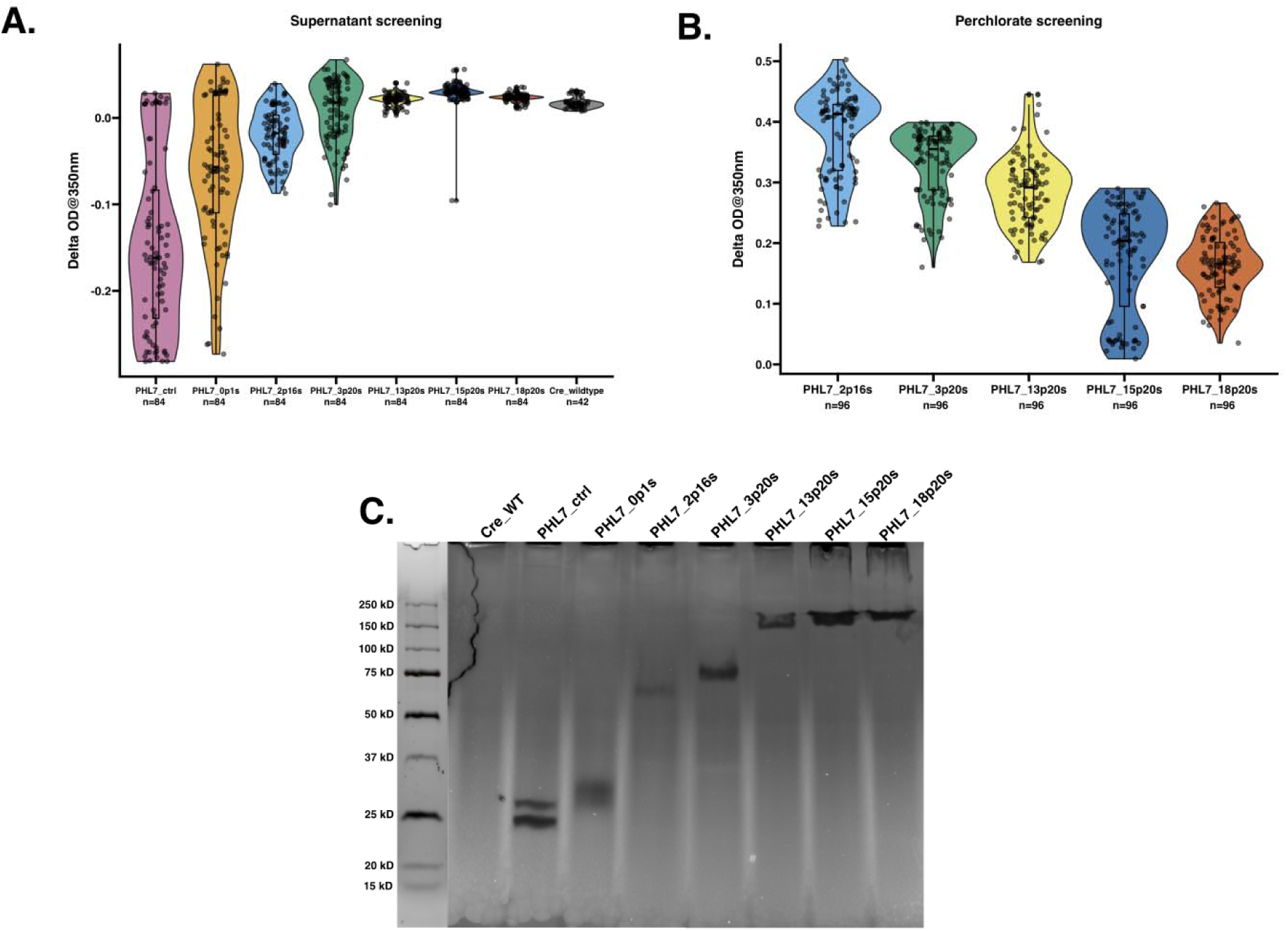
Screening of PHL7-glycomodule protein fusions for plastic-degrading activity using Impranil® as a substrate. **(A)** Violin plot showing the enzymatic activity of supernatants from *C. reinhardtii* transformants expressing PHL7 fused to GP1 fragments of varying lengths containing different numbers of PPSPX and SP motifs. Transformants were screened for plastic degradation using an Impranil® activity assay, measured as the change in optical density at 350 nm (ΔOD350). The data shows secretion of active protein into the supernatant from untreated cell cultures. Each dot represents the activity of an individual transformant. The violin plot illustrates the activity distribution, and the box plot indicates the interquartile range with a horizontal line marking the median activity for each construct. **(B)** Violin plot illustrating Impranil® DLN degradation activity of cell wall proteins released after perchlorate treatment. The presence of perchlorate in some experimental conditions alters the baseline optical properties of Impranil. Specifically, perchlorate can cause increased background absorbance over time due to colloidal destabilization. In the presence of active enzymatic degradation, this absorbance increase is counteracted, resulting in a net negative or reduced ΔOD350. Therefore, absolute OD values are less meaningful across different treatments, and comparisons are made within matched experimental sets. **(C)** Zymogram analysis of Impranil® degradation by secreted PHL7-glycomodule fusions. Clear bands indicate zones of enzymatic activity. Lanes correspond to wild-type *C. reinhardtii* (Cre_WT), a control strain expressing only PHL7 (PHL7_ctrl), and single transformants expressing various GP1 fusions. The molecular weight marker is shown in the most left lane. Note: Panels (A) and (B) collectively demonstrate the contribution of specific GP1 fragments to enzyme secretion and activity, while panel (C) confirms functional plastic degradation by the expressed PHL7-glycomodule fusions in a semi-quantitative gel-based assay.

Increasing the number of PPSPX motifs in GP1 fusion constructs led to a clear enhancement in anchoring efficiency, as demonstrated by reduced enzymatic activity in the supernatant (**Figure 2A**) and increased recovery following perchlorate treatment (**Figure 2B**). This trend was most evident among constructs that varied primarily in the number of PPSPX repeats while maintaining a relatively constant SP motif background (e.g., 16s to 20s). In contrast, the 0p1s construct, containing a single SP motif and lacking PPSPX repeats, showed significant secretion and minimal anchoring. These findings are consistent with prior reports that SP-rich glycomodules enhance secretion efficiency in both *C. reinhardtii* and *N. tabacum* [11–13], and further suggest that PPSPX motifs specifically promote cell wall retention.

Because diminishing activity was observed with the 2p16s, 3p20s, 13p20s, 15p20s, and 18p20s, these transformants were treated with 2 M sodium perchlorate to release the cell wall components. An Impranil® DLN-degrading activity assay was employed to screen for activity from PHL7 that could have been anchored to the cell wall via the glycomodule. The results now demonstrated a positive trend: as the number of motifs repeats increased, the activity levels also increased (**Figure 2B**). We observed statistically significant differences in activity between every glycomodule vector (ANOVA followed by Tukey post-hoc analysis; all had adjusted p-values < 0.0001) except for 15p20s and 18p20s (adjusted p-value 0.6578). The presence of perchlorate alters the baseline optical properties of Impranil® DLN. Perchlorate can cause increased background absorbance over time due to colloidal destabilization. In the presence of active enzymatic degradation, this absorbance increase is counteracted, resulting in a net negative or reduced ΔOD350. Therefore, absolute OD values are less meaningful across different treatments (**Figures 2A** and **B**), and comparisons are made within matched experimental sets.

### 2.2 Intein-mediated release of PHL7 from the cell wall to the extracellular environment

After pinpointing the potential anchor-signal sites for GP1, we investigated how to release PHL7 in a controlled manner. The PHL7-intein-GP1 vector was constructed, where the full-length GP1 and PHL7 coding sequences were fused via the pH intein [24]. The PHL7-intein-GP1 vector was transformed into *C. reinhardtii* via electroporation, and the cells were plated onto TAP agar plates containing 15 μg/mL zeocin and 0.75% (v/v) Impranil® DLN. The colonies with halos were picked into a 96-well plate containing TAP media, and grown for 7 days. An Impranil® DLN activity assay was performed to screen for positive transformants (**Supplementary Figure 1A**).

The transformant with the highest activity level was selected (here on referred to as W4_PHL7), and treated with pH 5.5 sodium acetate buffer (1 M NaCH_3_COO) or pH 8.0 potassium phosphate buffer (1 M K_2_HPO_4_). For W4_PHL7, the zymogram displayed only two clearing zones at about 24 kDa and 26 kDa for the pH 5.5 NaCH_3_COO-treated sample and no clearing zones for the pH 8.0 K_2_HPO_4_-treated sample (**Supplementary Figure 1B)**. We also observed a clearing zone at above 250 kDa (indicated by red arrow) and no clearing zones at 24 kDa and 26 kDa (indicated by blue arrow) for the perchlorate-treated sample, indicating an initially uncleaved form of the protein. These results exemplify that when the cells were exposed to pH 5.5 conditions, the pH intein successfully self-cleaved and released PHL7 from the cell wall.

### 2.3 Anchorage of mCherry to the cell wall via GP1

To investigate the functionality of the engineered intein in *C. reinhardtii*, we constructed a fusion protein consisting of mCherry linked to the intein and full-length GP1, referred to as mCherry-intein-GP1 (see **Plasmids** in https://doi.org/10.5281/zenodo.15823381). Screening of 84 transformants for mCherry fluorescence revealed that 28 colonies exhibited robust fluorescence, indicating successful expression of the fusion protein (**Figure 3A**). The control plasmid (mCherry-GP1), which lacks the intein, showed comparable levels of fluorescence. The wild-type (Cre_WT) showed minimal or no fluorescence.

**Figure 3:**
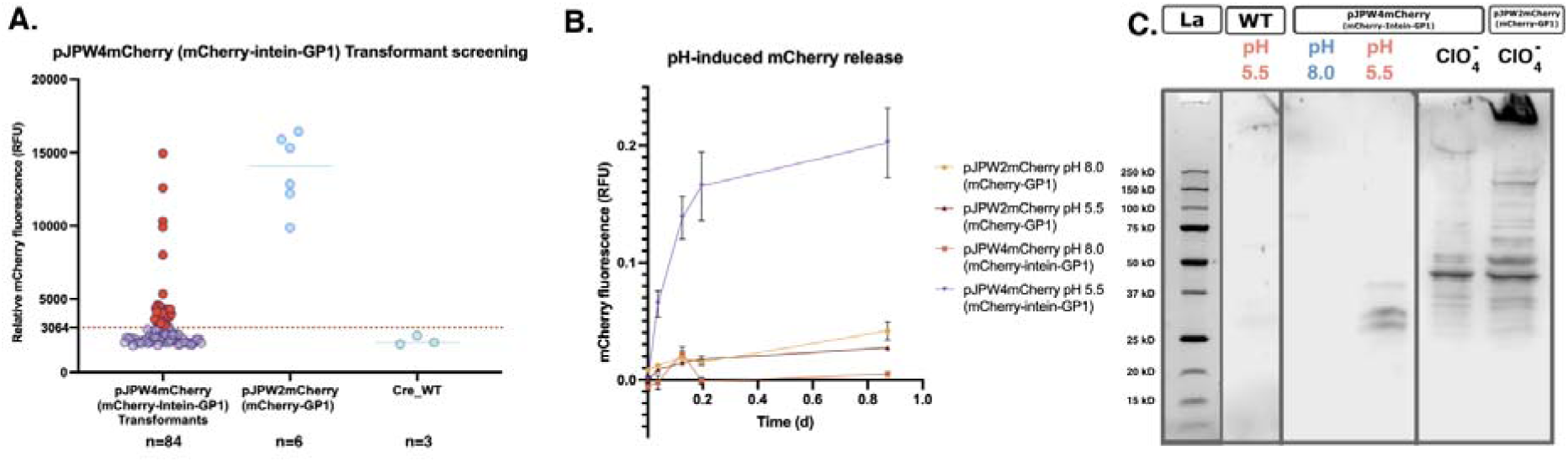
Analysis of mCherry release from C. reinhardtii recombinant strains unde different pH conditions and perchlorate treatment. **(A)** The mCherry-intein-GP1 vector was transformed into *C. reinhardtii*, and the transformant colonies were picked into 96-well plates containing TAP media. After 7 days of growth, the plates were screened for mCherry expression by measuring fluorescence at excitation wavelength 580/9 nm and emission wavelength of 610/20 nm. The threshold for positive expression of mCherry by the mCherry-intein-GP1 *C. reinhardtii* positive transformants was defined as three standard deviations above the mean fluorescence of the wild type (dotted line at 3064 RFU). A total of 28 transformants were above this threshold and are marked in red. **(B)** Time-course measurements of mCherry release from the top expressing mCherry-GP1 and mCherry-intein-GP1 strains under pH 5.5 and pH 8.0 conditions. The mean observed fluorescence values ± SD are plotted as a function of time (in days) are relative values from the total RFU observed after perchlorate treatment. The mCherry-intein-GP1 top expressor at pH 5.5 exhibited significantly higher release of mCherry compared to other conditions, suggesting enhanced release under acidic conditions. **(C)** Western blot analysis of mCherry protein in supernatants of the mCherry-intein-GP1 and mCherry-GP1 top expressors treated at different pH conditions and with perchlorate. Supernatant (left panel): mCherry was detected in the mCherry-intein-GP1 strain at pH 5.5 but not at pH 8.0, indicating pH-dependent release. Perchlorate treatment (right panel): Perchlorate-induced release of mCherry was observed for both the mCherry-intein-GP1 and mCherry-GP1 top expressors, with the latter displaying a most fully processed fusion protein.

To determine whether the intein underwent self-cleavage to release mCherry, transformants exhibiting the highest fluorescence from the initial screen were exposed to pH 8.0 and pH 5.5 conditions. Fluorescence kinetics demonstrated significant mCherry release from the mCherry-intein-GP1 top expressor at pH 5.5, with a measurable increase within 4 hours (**Figure 3B**). In contrast, release was minimal under pH 8.0 conditions and negligible for the mCherry-GP1 top expressor control, irrespective of the pH. These results indicate that the intein self-cleavage mechanism is strongly pH-dependent, favoring acidic conditions.

Western blot analysis confirmed the pH-dependent cleavage of the mCherry fusion protein (**Figure 3C**). At pH 5.5, mCherry was efficiently released from the mCherry-intein-GP1 top expressor, as shown by the presence of cleaved mCherry bands. Conversely, at pH 8.0, no cleavage products were detected. When cells were treated to release with perchlorate (ClO::J::J), distinct protein isoforms were observed for both mCherry-intein-GP1 and mCherry-GP1 top expressors possibly representing different glycosylation states of the fusion protein. Notably, the Western Blot shows an absence of the band at >250 kDa for the mCherry-intein-GP1 top expressor, which is otherwise present for the mCherry-GP1 top expressor. We hypothesize this to be the fully mature glycosylated form. We hypothesize thes absence of it in the mCherry-intein-GP1 top expressor could be due to altered glycosylation or processing, and future studies should investigate this.

To assess the potential impacts on cell viability and proliferation, growth curves of *C. reinhardtii* transformants expressing mCherry constructs (mCherry-intein-GP1 and mCherry-GP1) were compared to wild-type (Cre_WT). All strains displayed similar growth patterns over six days, as evidenced by overlapping 95% confidence intervals for the growth rate parameter, K (Gompertz model fit) across strains (**Figure 4A**). This indicates that neither the mCherry-intein-GP1 nor the control mCherry-GP1 constructs significantly affected cell growth. In parallel, mCherry fluorescence was monitored to evaluate protein expression levels over time (**Figure 4B**). The mCherry-intein-GP1 transformants exhibited lower fluorescence compared to mCherry-GP1, which lacks the intein, suggesting that intein-mediated cleavage or altered protein processing may reduce the accumulation of mCherry. This was also observed by the lack of overlapping 95% confidence intervals for production rate parameters (logistical curve fit) (see **Figure 4 Source Data** in https://doi.org/10.5281/zenodo.15823381). Wild-type cells showed negligible fluorescence, confirming the absence of background signal.

**Figure 4:**
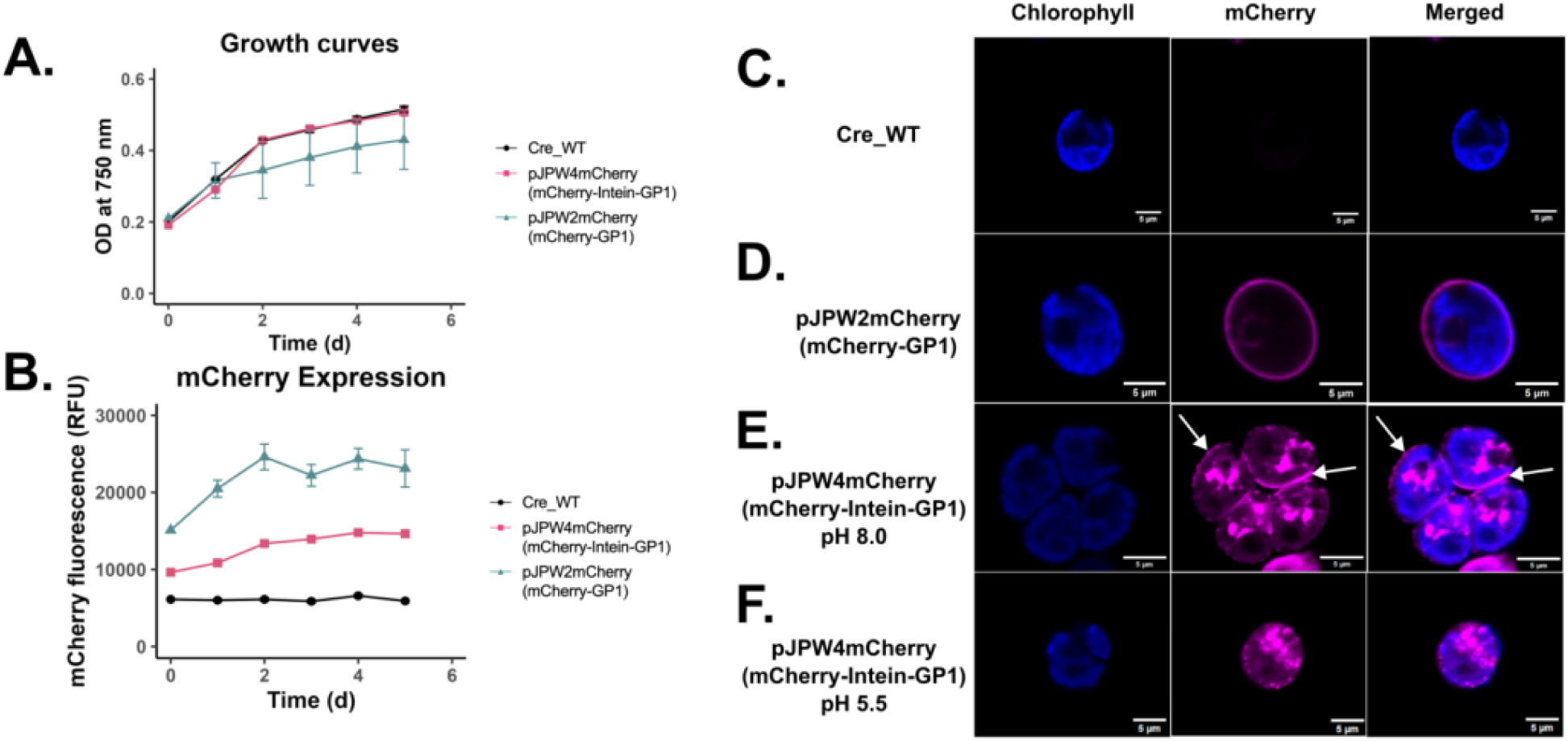
Growth analysis and microscopy visualization of C. reinhardtii strains expressing mCherry constructs, including pH-dependent localization effects. (A) Growth curves based on optical density at 750 nm over six days for wild-type (Cre_WT) and recombinant strains: mCherry-intein-GP1 (expressing mCherry::Intein::GP1), *pJP32mCherry* (secreting mCherry), and mCherry-GP1 (expressing mCherry::GP1). The growth dynamics of recombinant strains were similar to the wild type. This was also observed in the overlapping 95% confidence intervals for growth rate parameter, K (Gompertz model fit). Data points represent the mean ± SD from three biological replicates. **(B)** mCherry fluorescence measured in relative fluorescence units (RFU) over six days for the same strains. There was no overlap in the 95% confidence intervals for growth rate parameters (logistical curve fit). Recombinant strains exhibited higher mCherry fluorescence compared to the wild type. Data points represent the mean ± SD from three biological replicates. **(C)** Fluorescence microscopy images of wild type (Cre_WT) with no detectable mCherry fluorescence. **(D)** mCherry-GP1: Localization of mCherry::GP1 observed as a peripheral fluorescence pattern. **(E)** mCherry-intein-GP1 at pH 8.0: Aggregation or compartmentalization of mCherry observed, with fluorescence indicating retention within the cell and potentially part of the cell-wall (white arrows). **(F)** mCherry-intein-GP1 at pH 5.5: Altered fluorescence pattern with changes in mCherry localization or potential release from the cell wall under acidic conditions. Chlorophyll autofluorescence (blue) and mCherry fluorescence (magenta) are shown as individual and merged channels. All microscopy images include scale bars = 5 µm.

To further investigate the subcellular localization of mCherry, fluorescence microscopy was performed (**Figure 4C–F**). In the mCherry-GP1 top expressor (**Figure 4D**), mCherry fluorescence appeared consistently distributed around the cell periphery, likely localized to the cell wall [10]. However, in the mCherry-intein-GP1 top expressor (**Figure 4E**), mCherry signal was largely absent from the cell wall, with fluorescence observed predominantly in intracellular compartments (indicated by white arrows), likely corresponding to secretory vesicles. This pattern suggests incomplete secretion or improper targeting of the mCherry fusion protein in the presence of the intein. We did not observe a significant visual difference in mCherry location after pH treatment (**Figure 4E** and **F**). At pH 8.0, mCherry signal in mCherry-intein-GP1 cells remained intracellular, consistent with vesicular localization. Exposure to pH 5.5 resulted in slightly reduced mCherry signal from the cell wall. This observation may reflect pH-mediated cleavage and release of mCherry or experimental variability (e.g., sample preparation bias). Nevertheless, we observed a significant increase in released mCherry upon pH treatment (**Figure 3B**).

### 2.4 Investigating intein function through release of mCherry from the cell wall

To further characterize the behavior of the mCherry-intein-GP1 construct, we compared it to wild-type cells (WT, negative control), pJP32mCherry (a secreting strain), and mCherry-GP1 (a cell wall-localized strain). A nanofiltration-based fractionation assay was designed using a 100 kDa molecular weight cutoff filter, which allowed us to separate samples into retentate (≥100 kDa) and flowthrough (<100 kDa) fractions (**Supplementary Figure 2A**). This approach enabled the detection of mCherry fluorescence associated with either larger protein complexes or free cleaved mCherry.

The wild-type strain was used to account for any background fluorescence or autofluorescence caused by endogenous pigments. The pJP32mCherry strain was used as a control for mCherry secretion, while mCherry-GP1 served as a control for cell wall-associated mCherry signal. Experimental samples were subjected to pH 8.0 (neutral control) and pH 5.5 (cleavage-inducing condition), and fluorescence measurements were taken for both retentate and flowthrough fractions (**Supplementary Figure 2B**). mCherry fluorescence increased significantly in the pH 5.5-treated samples compared to pH 8.0 across all strains. In the WT strain, fluorescence increased by 2.21-fold in the retentate fraction (ANOVA, Tukey post hoc, p-value = 0.9228, n = 3 biological replicates), likely representing background signal unrelated to mCherry expression. For the pJP32mCherry strain, fluorescence increased 98.89 times in magnitude in the flowthrough fraction (p-value = 0.0006), consistent with secretion/leaking of mCherry. In the mCherry-GP1 top expressor, fluorescence increased by 10.00-fold in the retentate fraction (p-value < 0.0001), reflecting cell wall-associated mCherry. Fluorescence also increased 12.66 times in magnitude in the flowthrough fraction (p-value < 0.0001). This increase may be attributed to the secretion or leakage of partially glycosylated proteins (mCherry-GP1 fusion), which have a smaller molecular size, as observed in the Western blot (**Figure 3C**). Some mCherry proteins may be fully secreted without being anchored, contributing to the observed fluorescence signal. To account for this effect, fluorescence measurements of the culture supernatant are needed for normalization.

Interestingly, the mCherry-intein-GP1 top expressor exhibited a distinct fluorescence pattern, with significant increases observed in both the retentate (3.64-fold) and flowthrough (2.64-fold) fractions (p-values < 0.0001 and = 0.0001, respectively). These results suggest that the pH 5.5 treatment triggered mCherry cleavage and partial release from intracellular compartments and/or the cell wall. The concurrent fluorescence increases in both fractions further support the hypothesis that acidic conditions may cause partial cell lysis or destabilization (see Discussion for further analysis).

To investigate the possibility of cell lysis contributing to mCherry release under pH 5.5 conditions, we included an additional control strain, pAH04mCherry, which expresses mCherry cytosolically. All strains were treated at pH 8.0 and pH 5.5, and mCherry fluorescence in the supernatant was measured (**Supplementary Figure 3A**). Equal volumes of the supernatant from each strain were also analyzed using SDS-PAGE to visualize protein release patterns (**Supplementary Figure 3B**).

Across all strains, including the wild-type, an increase in mCherry signal was observed at pH 5.5 compared to pH 8.0 (**Supplementary Figure 3A**). For the wild-type strain, this increase was clearly detectable, reflecting background fluorescence likely originating from cell lysis and release of pigments (1.75 times higher mCherry RFU, ANOVA, Tukey, p-value < 0.0001, n = 3 biological replicates). In the cytosolic mCherry strain (pAH04mCherry), a significant increase in supernatant fluorescence was detected at pH 5.5 (4.48 times higher) compared to pH 8.0 (p-value <0.0001). This trend was similarly observed in the recombinant strains (pJP32mCherry, mCherry-GP1, and mCherry-intein-GP1), all of which exhibited elevated mCherry fluorescence at pH 5.5.

The SDS-PAGE analysis further corroborated these findings (**Supplementary Figure 3B**). At pH 5.5, the protein gel revealed a greater diversity of protein bands and overall higher intensity compared to pH 8.0, consistent with increased protein release under acidic conditions. Notably, the recombinant strains showed a more diverse protein release, further supporting the hypothesis of pH 5.5-induced cell membrane permeabilization.

To further characterize the role of pH and buffer composition in inducing cell lysis, we used the cytosolic mCherry strain (pAH04mCherry) to monitor the release of proteins under various buffer conditions. The tested conditions included a phosphate buffer (100 mM K_2_HPO::J, pH 8.0), a citrate buffer (100 mM trisodium citrate, pH 5.5), and acetate buffers at different concentrations (10 mM, 50 mM, 100 mM, and 200 mM sodium acetate, pH 5.5). mCherry fluorescence in the supernatant was measured as a proxy for cytoplasmic protein release (**Supplementary Figure 4**). The phosphate buffer at pH 8.0 resulted in a low background release of mCherry, likely due to background signal, or non-specific fluorescence values observed in the supernatant. In contrast, all acetate buffers at pH 5.5 caused a significant increase in mCherry fluorescence compared to the phosphate buffer, except for 50 mM acetate (ANOVA, Tukey post hoc, n = 3 biological replicates, see **Supplementary Figure 4 Source Data** in https://doi.org/10.5281/zenodo.15823381 for p-values).

Interestingly, the citrate buffer at pH 5.5 did not result in a significant increase in mCherry release compared to the phosphate buffer (p-value = 0.8871). This indicates that the effect of pH 5.5 is not solely due to acidity but is instead buffer-dependent, with acetate playing a specific role in promoting membrane destabilization or lysis. These results suggest that acetate buffer at pH 5.5 drives the release of cytoplasmic contents likely through a mechanism involving membrane permeabilization. In contrast, the lack of a similar effect with citrate buffer highlights a potential specificity in the interaction between acetate ions and cell membranes, which warrants further investigation.

## 3. Discussion

This study sheds light on key aspects of protein anchoring and controlled protein release in *C. reinhardtii*, with findings that have broad implications for biotechnology. First, we demonstrated that a specific fraction of the GP1 protein is necessary for anchoring of recombinant proteins to the cell wall. Using a series of truncated GP1 fragments fused to PHL7, we identified that the PPSPX motif plays a critical role in anchoring, despite containing the SP motif for secretion [11,12]. GP1, a widely studied glycoprotein, is known to reside in the chaotropic fraction of the cell wall and is composed of distinct structural domains, including a shaft region and a globular region [9,27]. Our results align with previous hypotheses suggesting that the PPSPX motif, which forms polyproline type II (PPII) helices, plays a crucial structural role in anchoring by mediating interactions with neighboring proteins [8]. Our results confirm that the PPSPX motifs are critical for anchoring proteins to the cell wall. Constructs with a higher number of PPSPX motifs displayed reduced secretion of PHL7 (pJP32PHL7_13p20s, pJP32PHL7_15p20s, and pJP32PHL7_18p20s), suggesting that these motifs promote retention within the cell wall (**Figure 2**). This is consistent with the proposed role of PPII helices in protein-protein interactions within the extracellular matrix.

Unexpectedly, the addition of GP1 sequences did not enhance secretion efficiency as some patents had suggested [12]. Other groups have previously demonstrated that the SP motifs associated with GP1 appear to facilitate enhanced protein secretion, as observed in both plants and algae [11–13]. Instead, our findings reveal that the PPSPX motif primarily acts as an anchoring domain, while the SP motifs likely influence efficient secretion. Ideally, a motif-by-motif dissection would involve variants containing exclusively one or the other motif class (e.g., 18p0s or 0p20s constructs), and we acknowledge that such constructs would provide stronger mechanistic clarity. However, the 20s construct has already been independently validated in previous studies [11–13], supporting the role of SP motifs in enhancing secretion. In our work, due to the high GC content and repetitive nature of the GP1 sequence, we were limited in the precise assembly of synthetic variants and instead relied on PCR-derived truncations that retained varying numbers of both motif types, more relevantly, varying the number of proline-rich PPSPX motifs to complement existing literature. Nevertheless, this new understanding clarifies the mechanistic roles of GP1’s distinct motifs and reveals the structural complexity of this glycoprotein in the cell wall.

The implications of these findings are both theoretical and practical. On a theoretical level, our results pinpoint PPSPX motifs as essential for cell wall anchoring, providing a foundation for understanding similar motifs in other *C. reinhardtii* proteins, such as those related to mating. For instance, the sexual agglutinins that are involved in sexual adhesion between gametes of *C. reinhardtii* contain a shaft domain in a PPII configuration that comprises PPSPX motifs [28]. These agglutinins are encoded by the *sag1* (sexual agglutination) and *sad1* (sexual adhesion) genes and mediate cell-cell interactions critical to the mating process. Furthermore, the PPSPX motifs forming PPII helices resemble those in structural proteins such as collagen, elastin, and actin-binding proteins [29], suggesting a conserved role across biological systems. This conservation implies that PPII helices contribute to stability and interaction dynamics in extracellular matrices and cytoskeletal systems across diverse organisms. This structure is also often found in the Src homology 3 (SH3) and tryptophan-tryptophan (WW) domains. The SH3 domain is involved in cell proliferation, migration and cytoskeletal modifications [30], and the WW domain is found in many signaling and structural proteins including RNA-binding proteins [31]. Moreover, the role of GP1’s C-terminal globular region, which was not directly addressed in this study, may warrant further investigation to determine how it contributes to cell wall integration and interactions with other matrix components.

From a practical perspective, understanding the distinct roles of the SP and PPSPX motifs could guide the development of constructs tailored for specific biotechnological needs. For instance, maximizing SP motifs could enhance protein secretion, while increasing PPSPX content could improve anchoring efficiency. Furthermore, different configurations could be strategically designed to optimize functionality. For example, anchoring domains could be positioned in the N-terminal region with SP motifs in the C-terminal region to facilitate both efficient secretion and anchoring of the engineered protein, or vice versa, depending on the specific application and structural requirements of the recombinant construct. Additionally, identifying PPSPX as a key anchoring motif offers opportunities for engineering smaller and more efficient anchoring modules. Such modules could be used to tether enzymes to surfaces for biocatalysis or to stabilize proteins within biofilms for industrial or environmental applications [32,33].

The experimental limitations of this study should also be considered. The truncated GP1 variants analyzed here were generated via PCR, which limited control over the exact motif composition. Future studies should employ synthetic constructs specifically designed to isolate the effects of SP and PPSPX motifs. This approach would pinpoint the minimal PPSPX motif content required for anchoring. While PCR artifacts were observed due to the GC-rich repeat regions, these do not reflect instability in *C. reinhardtii*, as it derives from the same endogenous sequence of the native GP1. Future work using long-read sequencing technologies could further verify structural integrity at the genomic level. Furthermore, the reliance on activity assays and zymograms, while informative, may not fully capture the structural dynamics of GP1 variants within the cell wall.

For many biotechnological applications, particularly those involving drug delivery or industrial bioprocessing, it is not sufficient to simply anchor proteins; there must also be a mechanism to release these proteins under specific conditions, enabling precise control over their activity. To address this need for controlled release, we successfully employed a cis-cleaving, pH-sensitive intein system, which complements the anchoring properties of glycomodules by enabling the conditional release of the anchored proteins under acidic conditions. The intein-mediated cleavage mechanism allowed for the release of PHL7 and mCherry under acidic conditions (pH 5.5), representing a novel approach to controlled release in microalgae. This advancement is particularly significant for applications requiring precise timing and spatial control, such as drug delivery systems. For instance, groups have previously engineered microalgae to carry payloads (e.g., antibiotics or small molecule cancer chemotherapeutic drugs) that get released via photocleavable linkers [16] or protease-cleavable linkers [34]. Moreover, for recombinant protein production, this system could simplify downstream processing by enabling the selective cleavage and harvesting of proteins in response to environmental stimuli, reducing the need for extensive purification steps.

Here, the cis-cleaving, pH-sensitive intein [24]functioned effectively in *C. reinhardtii*, facilitating the release of PHL7 and mCherry at pH 5.5. Zymogram analysis and fluorescence kinetics confirmed that cleavage is pH-dependent, favoring acidic environments while showing no activity at pH 8.0. This aligns with the known properties of the pH intein from Choi et al. and validates their application for tightly controlled protein release [24]. In the case of mCherry, microscopy revealed that intein-mediated cleavage also influenced protein processing and localization. The reduced presence of characteristic mCherry at the whole cell wall in mCherry-intein-GP1 transformants, combined with its prominent vesicular localization, suggests that the intein may interfere with proper protein trafficking or secretion. This was further corroborated by detecting several protein isoforms on a Western Blot which we hypothesize to be different glycosylation states. The full mCherry-intein-GP1 fusion protein is expected to have a molecular weight above 250 kD, as the native GP1 protein has a molecular weight of 272.4 kD when modified (and 65.1 kD when deglycosylated) [8]. Unmodified mCherry and the intein has a molecular weight of 26.7 kD and 18.6 kD, respectively. On the Western blot, we detected at least 22 bands to mCherry-GP1 and 17 bands to mCherry-intein-GP1 (**Figure 3C**), indicating isoforms potentially arising from variable glycosylation and sequential post-translational modifications. Notably, the fully processed protein observed in mCherry-GP1 was absent in mCherry-intein-GP1, implying that the intein might be disrupting parts of post-translational modifications. We hypothesize that these isoforms represent different glycosylation states, but glycan profiling should be conducted in future studies to investigate these isoforms.

Several limitations of the pH-sensitive intein must be addressed. The intein appeared to reduce overall mCherry accumulation compared to the control constructs, suggesting that intein presence in the tested configuration might affect protein yield. Localization and processing issues observed in mCherry transformants further indicate that intein integration can interfere with proper trafficking and secretion. The use of perchlorate revealed different isoforms on the Western Blot, which are possibly different glycosylation states that vary between constructs as discussed above. Furthermore, reliance on acidic conditions for cleavage may limit applicability in systems where pH sensitivity of host cells or proteins is a concern. Because acetate buffer at pH 5.5 induces release of cytosolic mCherry, this confounds the interpretation of cleavage-specific release. Direct testing of the mCherry-intein-GP1 construct in citrate buffer at pH 5.5 would provide a stronger mechanistic resolution, and should be the focus of future research. Our data suggest that citrate causes significantly lower release of intracellular mCherry than acetate across a range of concentrations, despite equivalent pH (**Supplementary Figure 4**). These findings suggest that buffer composition, in addition to pH, plays a critical role in modulating membrane integrity and potential cell lysis. While a portion of the mCherry release observed in **Figure 3B** may be attributed to lysis under acidic conditions, no detectable release, whether lytic or cleavage-mediated, was observed in the control mCherry-GP1 construct under the same conditions. Moreover, a distinct shift in banding pattern was observed in the Western Blot (**Figure 3C**), with the disappearance of the higher molecular weight fusion protein and the emergence of a band corresponding to free mCherry, consistent with intein-mediated cleavage..

To enhance the utility of intein-mediated systems, future research should focus on optimizing protein trafficking and processing by modifying intein sequences to minimize disruptions or by designing constructs with more efficient and compact anchoring domains. The observed disruption in trafficking could potentially stem from interference with the glycosylation machinery or spatial constraints within the tightly assembled cell wall matrix. Incorporating smaller anchoring domains could mitigate these issues by providing more space for proper protein anchoring thereby improving functional integration into the cell wall. Creating inteins with narrower pH sensitivity or tunable cleavage thresholds would allow for greater control in complex systems. Expanding the system to include other pH-sensitive inteins or engineering inteins responsive to additional stimuli, such as temperature or light, could further broaden its applications.

With insights into the GP1 glycomodules and intein-mediated controlled protein release, the observed pH-induced lysis in *C. reinhardtii* offers an additional layer of biological understanding and technological opportunity. Cell lysis, essential for applications such as protein purification, proteomics, and recombinant protein production, was effectively induced using acetate buffer at pH 5.5. This treatment resulted in the release of cell contents, including mCherry (**Supplementary Figure 4**). The mechanism underlying this phenomenon appears to involve the specificity of acetate in promoting lysis. While *C. reinhardtii* is known to be sensitive to acidic conditions [35], our findings suggest that the protonated form of acetate (i.e., acetic acid) may play an additional role as a surfactant, disrupting the cytoplasmic membrane. The specificity observed with acetate, compared to phosphate or citrate buffers at a similar pH, supports this hypothesis. Acetic acid’s pKa of 4.8 determines its protonation state at different pH levels. At pH 5.5, approximately 17% of acetate exists in the protonated form (CH::JCOOH), while the remaining 83% is present as acetate ions (CH::JCOO::J). This proportion of acetic acid (CH::JCOOH) may enhance its ability to integrate into and destabilize lipid bilayers, promoting membrane permeabilization. Unlike phosphate or citrate buffers at a similar pH, which did not induce lysis under similar conditions, acetate also uniquely enabled controlled permeabilization while leaving the chloroplast largely intact (data not shown). In **Supplementary Figure 4**, the 10mM acetate condition presented a higher variability, likely due to poor media removal in the washing steps, which is a potential technical limitation. This could have led to higher mCherry measurements for one of the samples, compared to the other buffers at higher acetate concentrations. Nonetheless, these findings highlight the utility of acetate buffers in facilitating cell lysis and open avenues for optimizing lysis conditions for various biotechnological applications. Additionally, we employed a Stain-Free SDS-PAGE gel (**Supplementary Figure 3B**), which uses a fluorescence-based protein detection method that is directly correlated with total protein load and distribution, similar to traditional Coomassie staining. This approach allowed us to detect global protein release patterns under different pH and buffer conditions. As shown, an increase in protein release was observed at lower acetate buffer pH.

The exact mechanism of acetate-induced lysis remains unclear and would require further exploration across a broader range of pH levels and buffer compositions, including mixtures and other weak acids such as propionic acid. Specific investigations into membrane lipid composition and interactions with protonated acetate could provide a mechanistic basis for the observed specificity. These experiments were beyond the scope of this work, which focused on demonstrating the phenomenon rather than pinpointing its underlying mechanism. Ultimately, this study establishes a framework for intein-mediated controlled protein release in *C. reinhardtii*, offering a versatile tool for synthetic biology and biotechnology while identifying key areas for refinement to enhance its broader applicability.

## 4. Conclusion

This study advances our understanding of protein anchoring, controlled protein release, and pH-induced lysis in *C. reinhardtii*, offering insights into algae biotechnology. We identified that the PPSPX motif in GP1 plays a crucial role in anchoring proteins to the cell wall, providing a foundation for engineering glycomodules tailored for diverse applications. Additionally, the successful deployment of a pH-sensitive intein system enabled precise temporal and environmental control over protein release, demonstrating its potential for applications in drug delivery, biocatalysis, and recombinant protein production. The observed pH-induced lysis, mediated by acetate buffer, highlights a cell permeabilization mechanism, paving the way towards innovative proteomics and protein purification approaches. Together, these findings establish *C. reinhardtii* as a versatile platform for synthetic biology and emphasize the potential for harnessing its biological systems to address critical challenges in industrial biotechnology.

## 5. Materials and Methods

### 5.1 Construction of Plasmids

All vectors were assembled using the pBlueScript II KS+ (pBSII) backbone and contain the nuclear promoter AR1 (HSP70/RBCS2). The *ble* gene was employed as a selection marker. For all vectors (except for pAH04mCherry), the gene of interest was fused to *ble* with a foot-and-mouth-disease-virus 2A self-cleavage peptide (FMDV-2A) and signal peptide (SP7). The pAH04mCherry vector does not contain SP7 because mCherry is targeted to the cytoplasm.

The pJP32PHL7 vector [25] served as a negative control for the glycomodule vectors, and contains the PHL7 sequence codon-optimized for *C. reinhardtii* that was purchased from IDT (Integrated DNA Technologies, San Diego, CA, USA). The pJP32PHL7_X glycomodule vectors (where X denotes the number of PPSPX and SP motifs) were constructed using the pJP32PHL7 vector as a backbone. These vectors include an additional 0p1s, 2p16s, 3p20s, 13p20s, 15p20s, and 18p20s sequences attached to the C-terminus of PHL7, where “p” stands for the PPSPX motif and “s” stands for the SP motif. These PPSPX- and SP-repeat sequences were prepared by PCR of the GP1 cell wall protein from the mCherry-GP1 (mCherry-GP1) vector [10] using forward primer 5’-CCTGGCGGTATCTTCAACTGCCCT-3’ and reverse primer 5’-CTTGACGGCCACGGGCG-3’. The variation in the number of serine-proline repeats resulted from unintended PCR products during amplification, due to the challenges posed by repetitive nature and the high GC content of the GP1 sequence. Consequently, we relied on PCR-derived truncations that retained varying numbers of both motif types.

The pJP32mCherry and pAH04mCherry vectors [36] served as controls for the pJPW4mCherry (mCherry-intein-GP1) and pJPW4PHL7 (PHL7-intein-GP1) intein vectors. Both vectors contain the mCherry sequence codon-optimized for *C. reinhardtii* that was purchased from IDT. The mCherry-GP1 vector [10] also served as a negative control for the intein vectors because it does not contain an intein. For mCherry-intein-GP1 and PHL7-intein-GP1, the incorporated intein was classified as a cis-cleaving pH-induced intein that self-cleaves at its C-terminal at pH 5.5 [24]. The intein was codon-optimized for *C. reinhardtii* and purchased from IDT. For mCherry-intein-GP1, the pJP32 backbone is used, and the fusion protein includes the full-length coding sequence of the endogenous GP1 protein, fused at its N-terminus to the pH-sensitive cis-cleaving intein and mCherry. Similarly, for PHL7-intein-GP1, the pJP32 backbone was also used, and the full-length coding sequencing of endogenous GP1 is fused at its N-terminus to the intein.

All vector pieces were assembled using NEBuilder® HiFi DNA Assembly (NEB - New England Biolabs) following the manufacturer’s protocol. All vectors contain restriction sites that flank the expression cassette (XbaI on the 5’ end and KpnI on the 3’ end) for linearization. The restriction enzymes were obtained from New England Biolabs (Ipswich, MA, USA). The final sequences can be found at ZENODO (https://doi.org/10.5281/zenodo.15823381.

### 5.2 C. reinhardtii Growth Conditions and Transformation

#### 5.2.1 Growth Conditions

The wildtype, cell wall-containing *C. reinhardtii* CC-1690 (mt+) strain was obtained from the Chlamydomonas Resource Center in St. Paul, MN, USA. The strain was propagated in TAP medium at 25 °C with constant illumination at 80 μmol photons m^-2^s^-1^ and agitated at 150 rpm on a rotary shaker.

#### 5.2.2. Transformation

The plasmids were doubly digested using XbaI and KpnI restriction enzymes (New England Biolabs, Ipswich, MA, USA), followed by purification with the Wizard SV Gel and PCR Clean-up System (Promega Corporation, Madison, WI, USA) without fragment separation. DNA concentration was quantified using the Qubit dsDNA High Sensitivity Kit (Thermo Fisher Scientific, Waltham, MA, USA). The digested plasmids were transformed into *C. reinhardtii* CC-1690 using electroporation. The cells were cultured to reach mid-logarithmic growth phase (3 x 10^6^ - 6 x 10^6^ cells/mL) in tris-acetate-phosphate (TAP) medium and maintained at 25°C with constant light exposure (80 μmol photons m^-2^s^-1^) on a rotary shaker at 150 rpm. The cells were concentrated by centrifugation at 3000 G for 10 minutes and resuspended to density of 3 x 10^8^ - 6 x 10^8^ cells/mL using MAX Efficiency™ Transformation Reagent for Algae (Catalog No. A24229, Thermo Fisher Scientific). For electroporation, 250 μL of the cell suspension and 500 ng of a vector plasmid (digested with XbaI and KpnI restriction enzymes) was chilled on ice for five to ten minutes in a 4-mm wide cuvette compatible with the Gene Pulser®/MicroPulser™ (BioRad, Hercules, CA). The cell-DNA mixture was electroporated using the GenePulser XCell™ device (BioRad, Hercules, CA) with a pulse of 2000 V/cm for 20 microseconds. Following electroporation, the cells were transferred into 10 mL of TAP medium and incubated with gentle agitation (50 rpm) in ambient light conditions (approximately 8 μmol photons m^-2^s^-1^) for an 18-hour recovery period. Post-recovery, the cells were concentrated by centrifugation, resuspended in 600 μL of TAP medium, and evenly spread onto two TAP agar plates with 15 μg/mL zeocin for mCherry-intein-GP1-transformed cells and two TAP agar plates with 15 μg/mL zeocin and 0.75% (v/v) Impranil® DLN for pJP32PHL7gXSP- and PHL7-intein-GP1-transformed cells. Impranil® DLN is a colloidal polyester-PUR dispersion [37]. The plates were incubated under a light intensity of 80 μmol photons m^-2^s^-1^ at 25 °C until visible colonies were formed. This transformation protocol can be found at [38].

### 5.3 Strain Screening

#### 5.3.1 mCherry Fluorescence Analysis

Transformant colonies were picked into 96-well plates: 84 transformant colonies, 6 wildtype *C. reinhardtii* CC-1690 colonies, and 6 blank wells. Each well contained 160 μL of TAP media.

After a growth period of 7 days, fluorescence measurements were conducted using the Infinite® M200 PRO plate reader (Tecan, Männedorf, Switzerland). Chlorophyll was measured at an excitation wavelength of 440/9 nm and emission wavelength of 680/20 nm for cell density normalization. mCherry fluorescence was measured at an excitation wavelength of 580/9 nm and emission wavelength of 610/20 nm for analyzing protein expression. Transformants exhibiting the highest mCherry-to-chlorophyll ratio were selected for further analysis (cell wall extraction, cell wall recrystallization, intein cleavage experiments, microscopy, and immunoblotting).

#### 5.3.2 Impranil® DLN Activity Assay

*C. reinhardtii* transformant colonies with clearing zones, or halos, were picked into two 96-well plates. Each plate contained 84 transformant colonies, 6 of the *C. reinhardtii* CC-1690 wild type colonies, and 6 blank wells. Each well contained 160 μL of TAP media. After a growth period of 7 days, fluorescence measurements were conducted using the Infinite® M200 PRO plate reader (Tecan, Männedorf, Switzerland). Chlorophyll was measured at an excitation wavelength of 440/9 nm and emission wavelength of 680/20 nm for cell density normalization. The 96-well plates were centrifuged at 2000 G for 5 minutes, and the supernatant was collected and filtered through a 0.2 µm PVDF Membrane 96-well filter plate (Corning®, Corning, NY, USA). The filtered supernatant (total soluble protein) was collected in a 96-well PCR plate (Fisher Scientific, Waltham, MA, USA) from which 50 µL was taken for the polyester polyurethane polymer degradation assay.

This protocol was adapted from Molino et al., 2024. The 96-well round bottom plate (Thermo Fisher Scientific, Waltham, MA, USA) was prepared by heating a mixture of 0.2% (w/v) agarose and 1 M K_2_HPO_4_ buffer solution in a 50 ml erlenmeyer flask in a microwave until the agarose is completely dissolved. Impranil® DLN was then added to the hot solution to reach a final concentration of 0.25% (v/v) and mixed thoroughly to ensure homogenous distribution. Subsequently, 150 μL of the molten Impranil® DLN-agarose solution was pipetted into each well of a 96-well round bottom plate and allowed to solidify at room temperature. Once the gel had solidified, 50 μL of the enzyme preparation was pipetted into each well. The assay plate was tightly sealed using a transparent 96 Well Plate Sealing Film (Lichen Cottage, Item Number SF-100). Using the Infinite® M200 PRO plate reader (Tecan, Männedorf, Switzerland), absorbance readings were measured at 350 nm, every 5 minutes, for over the course of 7 days. PHL7 activity was analyzed by calculating the change in absorbance over time. The presence of perchlorate in some experimental conditions alters the baseline optical properties of Impranil. Specifically, perchlorate can cause increased background absorbance over time due to colloidal destabilization. In the presence of active enzymatic degradation, this absorbance increase is counteracted, resulting in a net negative or reduced ΔOD350. Therefore, absolute OD values are less meaningful across different treatments, and comparisons are made within matched experimental sets.

#### 5.3.3 Zymograms with Impranil® DLN

Three biological replicates of the highest expressing clones from transgenic *C. reinhardtii* pJP32PHL7_X, pJP32PHL7, and the *C. reinhardtii* CC-1690 wild type were used for the zymogram. Strains were cultured for 5 days, and the 50 mL culture was split two ways: 1) supernatant collection and concentration and 2) chemical extraction of cell wall components. Supernatants containing total secreted soluble protein were collected from the transgenic *C. reinhardtii* pJP32PHL7_X, pJP32PHL7, and the *C. reinhardtii* CC-1690 wild type. These replicates were pooled together and each were equally concentrated 300X using 10 kDa centrifugal filters (Amicon® Ultra, Darmstadt, Germany). These samples were used for the first zymogram. To determine if PHL7 was anchoring to the cell wall, the chaotropic soluble portion of the transgenic *C. reinhardtii* pJP32PHL7_X cell wall was extracted, following the protocol adapted from [9]. Using the other half of the cultures, 1 mL was taken from each biological replicate and centrifuged at 3000 g for 2 minutes. The pellets were washed with 1 mL of ddH::JO, followed by centrifugation. The cell pellet was then resuspended in 150 μL of 2 M sodium perchlorate, centrifuged at 20,000 g for 1 minute, and 50 μL of the supernatant was used for the second zymogram.

The supernatant and perchlorate samples were separately mixed with 4X Laemmli Sample Buffer (BioRad, Hercules, CA). Equal volumetric amounts of protein samples (30 µL) were loaded into two 12% TGX Stain-Free™ FastCast™ gels containing 1% v/v Impranil® DLN (BioRad, Hercules, CA). The 12% TGX Stain-Free™ FastCast™ gels were prepared as per the manufacturer’s instructions, with the modification of adding 1% v/v Impranil® DLN to the solution to enable detection of PHL7 enzymatic activity. The proteins were separated at 80 V to 120 V. Following electrophoresis, the gels were immersed in a K_2_HPO_4_ buffer solution (100 mM, pH 8.0), and incubated at 37°C until clearing zones (i.e., transparent bands) appeared. This incubation step was crucial for the development of clearing zones, which indicates enzymatic degradation of the polyester polyurethane polymer within the gel matrix. Clearing zones typically emerged within 1-2 days of incubation, allowing for the qualitative assessment of enzymatic activity. This protocol was adapted from Molino et al., 2024.

#### 5.3.4 Cleavage of Intein with Sodium Acetate (pH 5.5) Treatment

##### 5.3.4.1 Screening for positive mCherry-intein-GP1 transformants

*C. reinhardtii* mCherry-intein-GP1 transformants were screened for mCherry fluorescence and the highest expressing strains were expanded to 50 mL cultures. *C. reinhardtii* pJP32mCherry, *C. reinhardtii* pAH04mCherry, and *C. reinhardtii* mCherry-GP1 strains were used as controls, and these strains were also expanded to 50 mL cultures. All strains were cultured for 5 days, and 1 mL of the culture was centrifuged at 3000 g for 2 minutes. The cell pellet was washed with 1 mL of ddH_2_O, followed by centrifugation at 3000 g for 2 minutes. The cell pellet was then resuspended in 1 mL of K_2_HPO_4_ (100 mM, pH 8.0) buffer or NaCH_3_COO (1 M, pH 5.5) buffer, and incubated at room temperature for 1 hour. After incubation, the cells were centrifuged at 20,000 g for 1 minute, and 100 μL of the supernatant was used for mCherry fluorescence reading at excitation wavelength of 580/9 nm and emission wavelength of 610/20 nm.

##### 5.3.4.2 Kinetics of pH-induced mCherry release from mCherry-intein-GP1 strain

The *C. reinhardtii* mCherry-intein-GP1, *C. reinhardtii* pJP32mCherry, *C. reinhardtii* pAH04mCherry, and *C. reinhardtii* mCherry-GP1 top expressors were used to measure mCherry release over time under pH 5.5 and pH 8.0 conditions. All strains were cultured for 5 days, and 1 mL of the culture was pipetted into 48-well plates (Corning®, Corning, NY, USA) in triplicates. The plate was centrifuged at 3000 g for 2 minutes, and the cell pellet was washed with 1 mL of ddH_2_O, followed by centrifugation at 3000 g for 2 minutes. The cell pellet was then resuspended in 1 mL of K_2_HPO_4_ (100 mM, pH 8.0) buffer or NaCH_3_COO (1 M, pH 5.5) buffer, and incubated at room temperature for 1 hour on a rotary shaker for gentle agitation (50 rpm). After incubation, the cells were centrifuged at 3000 g for 2 minutes, and 100 μL of the supernatant was used for mCherry fluorescence reading at excitation wavelength of 580/9 nm and emission wavelength of 610/20 nm. After each fluorescence reading, the 100 μL supernatant sample was put back into the 48-well plate to retain constant volume, and incubated at room temperature for 1 hour on a rotary shaker for agitation (50 rpm) again. This process was repeated 3 times, where fluorescence readings were taken every hour to track mCherry release over time.

#### 5.3.5 Cell membrane permeabilization using different buffers

The following buffers were prepared: 100 mM potassium dihydrogen phosphate at pH 8.0, 100 mM trisodium citrate at pH 5.5, and sodium acetate at pH 5.5 at concentrations of 10 mM, 50 mM, 100 mM, and 200 mM. A 50 mL culture of the pAH04mCherry strain expressing cytosolic mCherry was grown for 5 days. In three replicates, 6 x 1 mL of the culture was centrifuged at 3000 g for 2 minutes. The cell pellets were washed with 1 mL of ddH_2_O, followed by centrifugation at 3000 g for 2 minutes. The cell pellets were then resuspended in 1 mL of the different buffers, and incubated at room temperature for 1 hour. After incubation, the cells were centrifuged at 20,000 g for 1 minute, and 100 uL of the supernatant was used for mCherry fluorescence reading at excitation wavelength of 580/9 nm and emission wavelength of 610/20 nm.

### 5.4 Chemical Extraction of Cell Wall Components and Recrystallization of mCherry

This protocol was adapted from Goodenough et al. to extract the chaotropic soluble portion of the *C. reinhardtii* cell wall [9]. Strains were cultured for 5 days, and 1 mL of the culture was centrifuged at 3000 g for 2 minutes. The pellet was washed with 1 mL of ddH::JO, followed by centrifugation. The cell pellet was then resuspended in 150 μL of 2 M sodium perchlorate, centrifuged at 20000 g for 1 minute, and 100 μL of the supernatant was used for fluorescence reading at excitation wavelength of 580/9 nm and emission wavelength of 610/20 nm.

For cell wall recrystallization, 40 mL of culture was centrifuged at 3000 g for 3 minutes. The pellet was washed with 40 mL of ddH::JO and resuspended in 1 mL of 2 M sodium perchlorate. After transferring to a microcentrifuge tube, the mixture was centrifuged at 20000 g for 1 minute. Approximately 900 μL of the supernatant was concentrated to 50 μL using a 30 K centrifugal filter (Amicon® Ultra, Darmstadt, Germany), followed by three diafiltration steps with 450 μL ddH::JO. The cell wall crystals were recovered according to the manufacturer’s instructions by inverting the filter membrane into a new collection tube.

### 5.5 Cellular Fluorescence Localization

Transformed strains were cultivated in TAP medium until they reached the late log phase at 25°C, under continuous illumination of 80 μmol photons/m²/s, with agitation at 150 rpm on a rotary shaker. Live cells were then observed using agarose pads, prepared according to the protocol described at dx.doi.org/10.17504/protocols.io.bkn8kvhw. These cells were placed onto TAP 1% agarose pads, prepared with Frame-Seal™ Slide Chambers (15 × 15 mm, 65 μL) on a glass slide and covered with a coverslip before image acquisition. Live-cell imaging was conducted using an automated inverted confocal laser scanning microscope (Leica Stellaris 5 Confocal). mCherry fluorescence was excited at 580 nm with a laser set to 5% power, and emission was detected using a HyD hybrid detector set between 601 nm and 634 nm. Chlorophyll fluorescence was excited at 405 nm with a laser set to 2% power, and emission was detected between 650 nm and 750 nm, again using the HyD hybrid detector. Microscope settings were maintained consistently across all imaging sets, and images were acquired in sequence mode for both chlorophyll and mCherry. Image analysis was performed using Fiji, an ImageJ distribution, with uniform settings applied across all image groups. Brightness adjustments were made in Fiji, with consistent settings used unless otherwise noted. For video imaging, a Zeiss Elyra 7 Lattice SIM microscope was used. In this setup, a diode-pumped solid-state laser with a 561 nm wavelength was used to excite mCherry, with BP 525/50 and BP 617/73 filters in line with a long-pass dichroic mirror SBS LP 560 to observe mCherry fluorescence.

### 5.6 Western Blot

After mCherry fluorescence screening of *C. reinhardtii* pAH04mCherry- and mCherry-intein-GP1-transformants, the highest expressing strains were expanded to 50 mL cultures, along with the *C. reinhardtii* wild type. For the pAH04mCherry strain, the 50 mL culture was split in half, each of the 25 mL was centrifuged at 3000 g for 10 minutes, and the pelleted cells were treated with 1 mL (equal volume) K_2_HPO_4_ buffer (100 mM, pH 8.0) or NaCH_3_COO buffer (100 mM, pH 5.5) for 2 hours at room temperature under static condition. After incubation, the mixture was centrifuged at 20,000 g for 1 minute. The supernatants were collected and concentrated approximately 66.67X using 10 kDa centrifugal filters (Amicon® Ultra, Darmstadt, Germany). For the mCherry-intein-GP1 top expressor, the 50 mL culture was split three ways, two of which were used for K2HPO4 pH 8.0 buffer and NaCH_3_COO pH 5.5 buffer treatment, while the last aliquot was used for 2 M NaClO_4_ buffer treatment. The mCherry-GP1 top expressor was also expanded to 50 mL, and prepared for 2 M NaClO_4_ buffer treatment. The 15 mL cultures were centrifuged at 3000 g for 10 minutes, the pellet was washed with 15 mL of ddH::JO and resuspended in 1 mL of 2 M sodium perchlorate. After transferring to a microcentrifuge tube, the mixture was centrifuged at 20,000 g for 1 minute. The supernatants were collected and used for the Western blot.

All samples were denatured by adding 4X Laemmli Sample Buffer (BioRad, Hercules, CA), followed by boiling at 98°C for 10 minutes. Equal volumetric amounts of protein samples (30 µL) were loaded into the 12% TGX Stain-Free™ FastCast™ gel (BioRad, Hercules, CA). The proteins were separated at 80 V to 120 V. After separation, the proteins were then transferred to a nitrocellulose membrane at 15 V for 1 hour. The membrane was blocked with 2% Bovine Serum Albumin (Sigma-Aldrich, St. Louis, MO, USA) diluted in 1X phosphate-buffered saline with Tween® detergent overnight at 4°C on a rocking shaker (BioRocker™ 2D Rockers, Thermo Fisher Scientific, Waltham, MA, USA). After blocking, the membrane was probed with an anti-RFP polyclonal antibody conjugated to horse radish peroxidase (Abcam, Cambridge, United Kingdom) for 1 hour at room temperature on a rocking shaker (BioRocker™ 2D Rockers, Thermo Fisher Scientific, Waltham, MA, USA). Bands were detected using Clarity Western ECL Blotting Substrate (BioRad, Hercules, CA) as per the manufacturer’s instructions.

### 5.7 Separation of Proteins Using 100 kDa Membrane Filters After Cleavage of Intein With Sodium Acetate (pH 5.5) Treatment

After mCherry fluorescence screening of *C. reinhardtii* mCherry-intein-GP1-transformants, the highest expressing strains were expanded to 50 mL cultures. The highest expressing *C. reinhardtii* pJP32mCherry and *C. reinhardtii* mCherry-GP1 strains were used as controls, and these strains were also expanded to 50 mL cultures. All strains were grown for 5 days and the cultures were split in half for perchlorate treatment (1M) and sodium acetate (100 mM, pH 5.5) treatment. For each treatment group, 1 mL of the culture were centrifuged at 3000 g for 2 minutes, and the pellets were washed with K_2_HPO_4_ buffer (100 mM, pH 8.0), followed by centrifugation at 3000 g for 2 minutes. Then, the pellets were resuspended with 1 mL of either 1 M NaClO_4_ buffer or 100 mM NaCH_3_COO (pH 5.5) buffer. The cells were incubated at room temperature for 1 hour on a rotary shaker for agitation (50 rpm). After incubation, the cells were centrifuged at 20,000 g for 1 minute, and the supernatants were transferred into a 100 kDa centrifugal filter (Amicon® Ultra, Darmstadt, Germany) to centrifuge at 10000 g for 30 minutes. After centrifugation, the retentate and permeate samples were transferred to a 96-well plate and read for mCherry fluorescence at excitation wavelength of 580/9 nm and emission wavelength of 610/20 nm.

### 5.8 Data Analysis

R Statistic version 3.6.3 (2020-02-29) running in the RStudio v1.2.5042 IDE was used to import and generate plots for the figures and the statistical summary. The ggprism theme was used to generate graphs and plots [39]. GraphPad Prism 7.0 was also used to generate graphs and plots and perform statistical analyses. All data were analyzed and processed using Microsoft Excel 365 (Version 16.0, Microsoft Corporation, Redmond, WA, USA).

## Data Availability

The datasets generated during and/or analyzed during the current study are available in the ZENODO repository: https://doi.org/10.5281/zenodo.15823381

## Acknowledgements

The authors gratefully acknowledge the support of imaging specialist Marcy Erb, Ph.D. for her contribution in the microscopy core.

## Author contributions: CRediT

**KK:** Investigation, Methodology, Formal analysis, Visualization, Writing – original draft, review & editing

**EES:** Investigation, Methodology, Writing – review & editing

**CJD:** Investigation, Writing – review & editing

**SM:** contributed to drafting and revising the original manuscript and secured funding for the research.

**JVDM**: Conceptualization, Data curation, Formal analysis, Investigation, Methodology, Visualization, Writing – original draft, Writing – review & editing

## Supplementary Figures

**Supplementary Figure 1:**
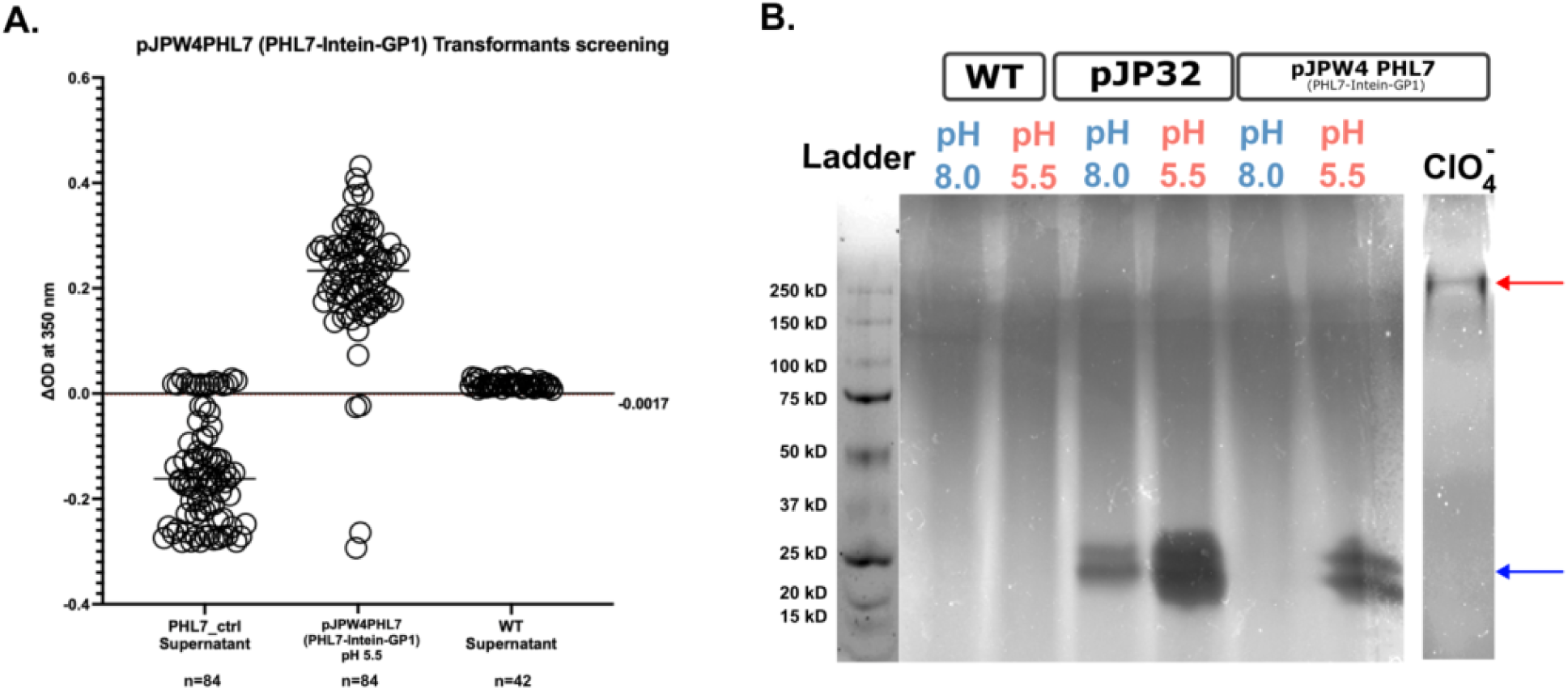
Zymogram analysis of plastic-degrading activity in *C.reinhardtii* strains expressing PHL7 under different pH conditions and perchlorate treatment. **(A)** Impranil® activity assay to screen for positive PHL7-intein-GP1 transformants. The threshold (red dotted line) for positive transformants represents three standard deviations below the mean WT supernatant activity (−0.0017). **(B)** This zymogram, performed using Impranil®, a plastic dispersion substrate, detects plastic degrading activity of the plastic-degrading enzyme PHL7 in samples collected from wild type (*WT*), pJP32 (PHL7-secreting top expressing strain), and PHL7-intein-GP1 (PHL7-intein-GP1 fusion protein-top expressing strain). The WT strain exhibited no detectable activity at either pH 8.0 or pH 5.5. The pJP32 strain, which secretes PHL7, showed clear plastic degrading activity at both pH 8.0 and pH 5.5, indicating functional secretion of the plastic-degrading enzyme. In contrast, the PHL7-intein-GP1 top expressor displayed no detectable activity at pH 8.0, but strong plastic degrading activity was observed at pH 5.5, consistent with pH-dependent cleavage and release of the active enzyme from the GP1-Intein-PHL7 fusion construct. Additionally, treatment of the PHL7-intein-GP1 strain with perchlorate (ClO) resulted in a clearing zone at above 250 kD (red arrow), indicating an initially uncleaved form of the protein, in contrast to the expected ∼25 kDa for the free form (blue arrow). This result highlights the pH- and chemical-dependent regulation of PHL7 release and activity in engineered *C. reinhardtii* strains.

**Supplementary Figure 2:**
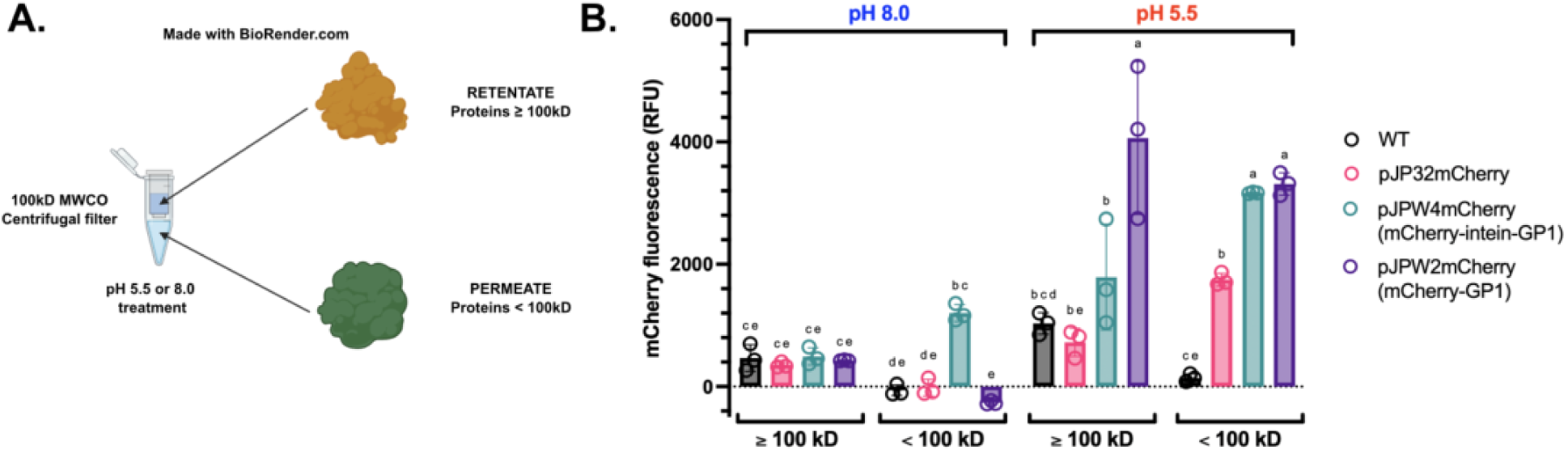
Analysis of mCherry fluorescence in supernatants of *C. reinhardtii* strains treated at pH 8.0 and pH 5.5 and filtered through a 100 kDa molecula weight cutoff (MWCO) filter. **(A)** Schematic showing a nanofiltration-based fractionation assay using a 100 kDa molecular weight cutoff filter (MWCO), which allowed us to separate samples into retentate (≥100 kDa) and flowthrough (<100 kDa) fractions. Made with BioRender.com. **(B)** The bar plots show mCherry fluorescence, measured in relative fluorescence units (RFU), in two filtrate fractions: proteins smaller than 100 kDa (“<100 kDa”) and retained proteins larger than 100 kDa (“>100 kDa”). Samples were analyzed from wild type (WT), pJP32 (secreting non-fused mCherry), mCherry-GP1 (top expressor), and mCherry-intein-GP1 (top expressor). **pH 8.0 (left panel)**: Low fluorescence was detected in wild-type, pJP32 *and* mCherry-GP1 fractions across both MWCO filters fractions. mCherry-intein-GP1 strains demonstrated higher fluorescence in the <100 kDa fraction, suggesting partial release of mCherry-containing proteins under neutral conditions. **pH 5.5 (right panel):** Acidic treatment resulted in a substantial increase in fluorescence in the <100 kDa fraction for all recombinant strains, pJP32, mCherry-GP1 and mCherry-intein-GP1, suggesting pH-dependent release of mCherry. pJP32 exhibited modest fluorescence in the <100 kDa fraction, consistent with secretion of mCherry. The >100 kDa fraction fluorescence decreased significantly, particularly for mCherry-intein-GP1, indicating potential dissociation or cleavage of the fusion protein under acidic conditions. Letters above bars indicate statistical groups, with distinct letters denoting significant differences (p < 0.05).

**Supplementary Figure 3:**
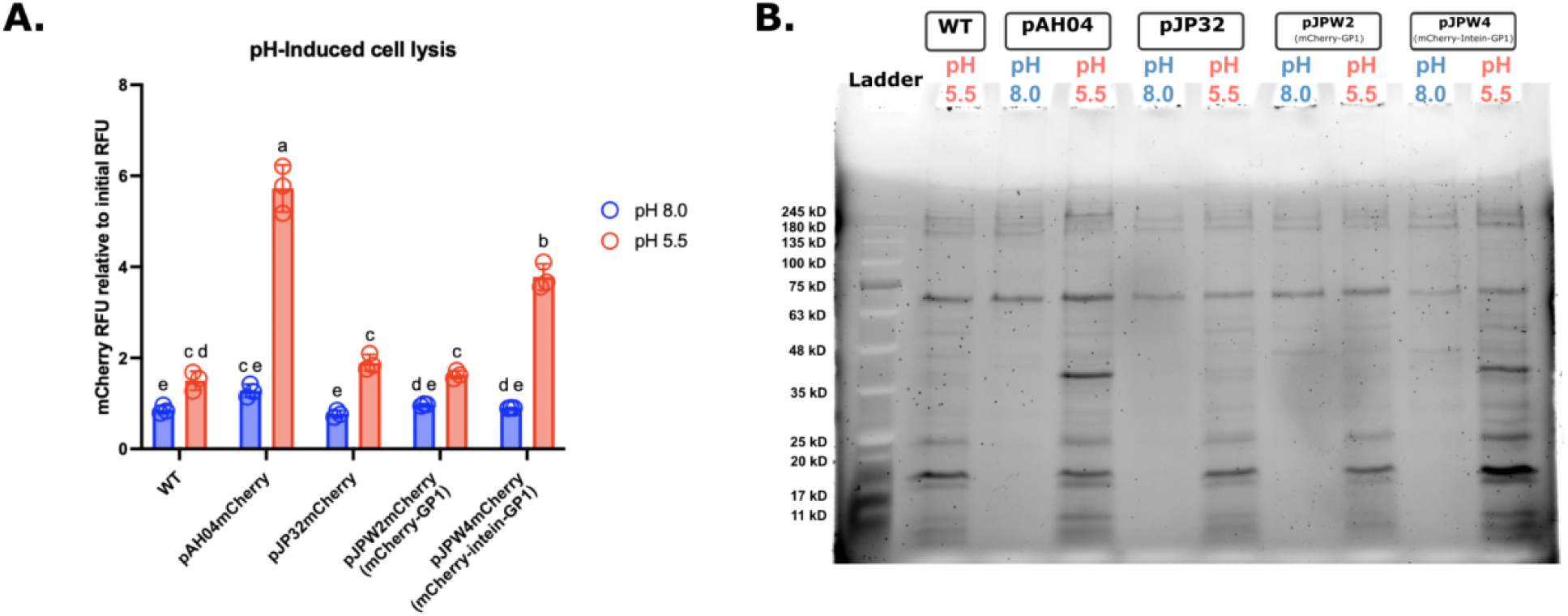
Analysis of pH-induced cell lysis in *C. reinhardtii* strains expressing mCherry. **(A)** Quantification of mCherry fluorescence released into the supernatant over 2 hours for the strains measured at excitation/emission 580/610 nm. Fluorescence values for the 2-hour time point were normalized to time-zero values and are shown for both pH 8.0 and pH 5.5. Error bars represent the standard deviation of 3 biological replicates. Strains expressing cytosolic mCherry (pAH04mCherry) showed significantly higher mCherry fluorescence release at pH 5.5 compared to pH 8.0, consistent with pH-induced cell lysis. Significant differences between groups are indicated by different letters (e.g., a, b, c, d; P < 0.05). **(B)** Stain-free SDS-PAGE gel showing total protein content in the supernatants of *C. reinhardtii* strains cultured in 100mM phosphate buffer at pH 8.0 or 100 mM acetate buffer at pH 5.5 for 2 hours. The lanes include a molecular weight ladder (Ladder) and the following strains: wild type (WT), pAH04mCherry (cytosolic expression of mCherry), pJP32mCherry (secretion of mCherry), mCherry-GP1 (top expressor), and mCherry-intein-GP1 (top expressor). Supernatants from both pH conditions were analyzed for all strains except the WT, which is shown only at pH 5.5 as a control.

**Supplementary Figure 4:**
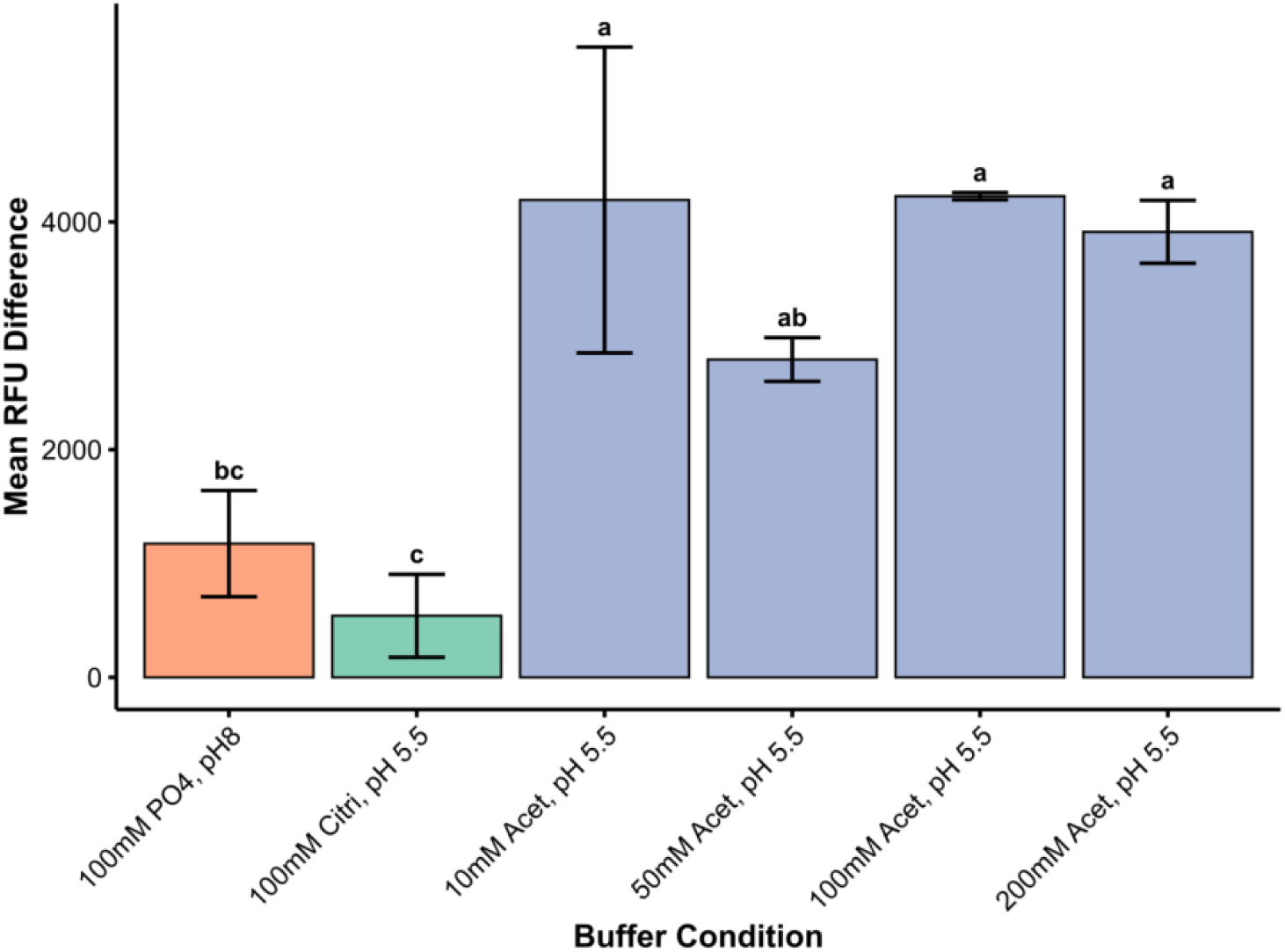
Buffer type, pH, and concentration influence cell lysis in *C. reinhardtii* expressing cytosolic mCherry. Mean mCherry fluorescence release (RFU difference) from the supernatant of *C. reinhardtii* strains expressing cytosolic mCherry after 4 hours under different buffer conditions. Buffers include 100 mM phosphate (PO[) at pH 8.0, 100 mM citrate (Cit) at pH 5.5, and acetate (Acet) at pH 5.5 at concentrations of 10 mM, 50 mM, 100 mM, and 200 mM. Fluorescence readings (Delta RFU) represent mCherry release due to cell lysis, normalized to initial time-zero values. Statistical groupings (letters: a, b, c, etc.) were determined using one-way ANOVA followed by Tukey’s post hoc test with three biological replicates per treatment (P < 0.05). Error bars represent the standard deviation of replicates.

